# Marmoset Model for *Mycobacterium avium* Complex Pulmonary Disease

**DOI:** 10.1101/2021.11.15.468600

**Authors:** Jay I Peters, Diego Jose Maselli, Mandeep Mangat, Jacqueline J. Coalson, Ceci Hinojosa, Luis Giavedoni, Barbara A. Brown-Elliott, Edward D. Chan, David E. Griffith

## Abstract

**Rationale:** *Mycobacterium avium* complex, is the most common nontuberculous mycobacterial respiratory pathogen in humans. Disease mechanisms are poorly understood due to the absence of a reliable animal model for *M. avium* complex pulmonary disease.

**Objectives:** Assess the susceptibility, immunologic and histopathologic responses of the common marmoset (*Callithrix jacchus*) to *M. avium* complex pulmonary infection.

**Methods:** 7 adult female marmosets underwent endobronchial inoculation with 10^8^ colony-forming units *of M. intracellulare* and were monitored for 30 or 60 days. Prior to infection, chest radiograph and serum cytokines were assessed; serum cytokines were also monitored weekly for 30 days. At sacrifice 30 days (3 animals) or 60 days (4 animals) after infection, chest radiograph, serum and bronchoalveolar lavage cytokines, histopathology, and cultures of the bronchoalveolar lavage, lungs, liver, and kidney were analyzed.

**Measurements and Main Results:** Five of seven animals (two at 30 days and three at 60 days of infection) had positive lung cultures for *M. intracellulare*. Extra-pulmonary cultures were positive in three animals. All animals appeared healthy throughout the study. All five animals with positive lung cultures had radiographic changes consistent with pneumonitis. At 30 days, those with *M. intracellulare* lung infection showed granulomatous inflammation while at 60 days there was less inflammatory change, but bronchiectasis was noted. The cytokine response in the bronchoalveolar lavage fluid was uniformly greater in the animals with positive *M. intracellulare* cultures than those without a productive infection with greater levels at 30-days compared to 60-days. Similarly, serum cytokines were more elevated in the animals that had positive *M. intracellulare* cultures compared to those without a productive infection, peaking 14-21 days after inoculation.

**Conclusion:** Endobronchial instillation of *M. intracellulare* resulted in pulmonary mycobacterial infection in marmosets with a differential immune response, radiographic and histopathologic abnormalities, and an indolent course consistent with *M. avium* complex lung infection in humans.

## INTRODUCTION

*Mycobacterium avium* complex (MAC) is the most common nontuberculous mycobacterial (NTM) respiratory pathogen in humans (1,2). MAC comprises multiple species and subspecies including *M. avium* and *M. intracellulare*, the two most important MAC respiratory pathogens (3-5). Typically, these two species are both reported as “MAC,” but their environmental sources differ, and there is evidence indicating differential pathogenicity and clinical disease severity between the two species (6,7). Since MAC is ubiquitous in the environment and exposure is likely unavoidable, it is apparent that some form of host susceptibility must also be present for MAC lung disease to occur (3,4,8). In that context, pulmonary MAC disease occurs primarily in patients with structural lung disease, especially bronchiectasis and emphysema, without demonstrable systemic immune suppression (3,4,8). Pathophysiologic questions are further complicated because MAC lung disease can evolve in two forms, either fibro-cavitary disease similarly to pulmonary tuberculosis (TB), or as a more indolent infection associated radiographically with nodules and bronchiectasis (nodular/bronchiectatic disease) (3-5).

A major impediment to greater understanding of fundamental pathophysiologic mechanisms surrounding MAC lung disease is the lack of a reproducible animal model that can replicate cellular, biochemical and pathological events observed in human MAC lung disease. The relevance of murine models of MAC infection for human MAC lung disease is not established and has uncertain applicability (9). Presumably, MAC lung infection in a host species more related to humans would be more informative and has been achieved with endobronchial instillation of *M. avium* in a rhesus macaque (10). While it also remains unclear if the mechanisms of disease establishment and progression in this model are pertinent to human MAC lung disease (10), an animal model using a species phylogenetically even closer to humans should provide a better approximation to the human mycobacterial response.

The common marmoset (*Callithrix jacchus*) has been used as a model for TB lung infection (9-12). Endotracheal instillation of TB strains in marmosets results in pulmonary abnormalities covering the entire spectrum of lesions observed in human TB patients including cavitation (11-14). However, the utility of marmosets as a model for human TB disease is limited because marmosets are exceptionally susceptible to *M. tuberculosis* (11-14). This susceptibility to mycobacterial infection combined with pathophysiologic similarities between marmosets and humans to *M. tuberculosis* infection are potential advantages for utilizing marmosets to study infection with a relatively non-virulent mycobacterial pathogen for humans such as MAC. Additionally, marmosets are smaller than macaques with relatively shorter life span, allowing feasible studies into advanced age. For these reasons, we investigated the potential of marmosets as a model for MAC lung disease.

## METHODS

### Non-human primate research regulations

Seven adult colony-bred female marmosets (*Callithrix jacchus*) were purchased from the Southwest National Primate Center, San Antonio, TX, and were found to be free of known primate bacterial and viral pathogens based on routine surveillance. The study and use of non-human primates were conducted in accordance with the Guidelines established by the Weatherall report and conformed to National Institutes of Health guidelines (15, 16). All animal work was approved by the University of Texas Health Science Center, San Antonio (UTHSC-SA) and the University of Texas Health Science Center, Tyler (UTHSCT) Institutional Animal Care and Use Committees and the Southwest National Primate Research Center Institutional Animal Care and Use Committee of the Texas Biomedical Research Institute (Texas Biomed). Animals were housed separately and had vital signs (heart rate, temperature and oxygen saturation) monitored closely throughout the experimental period. They were weighed daily, measured for nutritional and fluid intake, and examined twice daily for normal interactions with staff members.

### Animal Infection and Sample Collection

The seven adult marmosets were inoculated endobronchially at the level of the main carina using a special narrow diameter bronchoscope with one mL of a 10^8^ CFU/mL *M. intracellulare* obtained from the Mycobacteria/Nocardia Research Laboratory at the UTHSCT. All procedures (bronchoscopy, blood draws and euthanasia) were conducted under ketamine anesthesia with the additional use of isoflurane anesthesia with bronchoscopy and bronchoalveolar lavage (BAL) in the presence of veterinary staff. Each animal underwent assessment of serum chemistry, and complete blood count prior to inoculation and on the day of euthanasia. Because there are no previous comparable studies with this primate, we sacrificed a group of animals at 30 days and another group at 60 days to optimize the chance of recovering *M. intracelluare* as well as to define the time course of an evolving inflammatory response. Cytokine analysis was obtained prior to inoculation with *M. intracellualre* and on a weekly basis from day 0 to day 30 for all animals and again on day 60 for the animals sacrificed at day 60. All the animals had BAL performed prior to euthanasia at either 30- or 60-days post-inoculation. The animals were then taken directly to necropsy by a primate pathologist.

### Imaging

Supine postero-anterior and lateral chest X-rays were obtained at baseline and prior to sacrifice.

### Histopathology

Formalin-fixed, paraffin-embedded tissue sections were deparaffinized and stained with hematoxylin and eosin for histopathological analysis as well as for acid-fast bacteria (AFB) stain. Histopathologic images were obtained using an Axioplan microscope (Carl Zeiss, Jena, Germany) with a Spot Insight camera (Diagnostic Instruments Inc., Sterling Heights, MI.)

### Microbiologic Assessments

A macrolide- and aminoglycoside-resistant (clarithromycin MIC >16 μg/mL and amikacin MIC >64 μg/mL) isolate of *M. intracellulare*, previously identified by 16S rRNA gene sequence, was prepared for inoculating the marmosets. The clinical isolate was grown on Middlebrook 7H10 agar. After adequate growth was obtained (approximately 7-10 days), several colonies were transferred to 3 mL of sterile distilled water to prepare a suspension with optical density equal to a 0.5 McFarland standard by nephelometer reading. The inoculum was chosen since this turbidity represents the approximate number of organisms (10^8^ CFU/mL) present in the matched turbidity McFarland standard used for antimicrobial susceptibility testing as recommended by the Clinical and Laboratory Standards Institute (CLSI) (17). The suspension was incubated for 7 days at 35°C and 1-3 mL aliquots prepared to be used to inoculate the marmosets.

BAL and tissue samples were processed and cultured for mycobacteria by the Mycobacteria/Nocardia Research Laboratory at the UTHSCT, using standard decontamination procedures, fluorochrome microscopy, solid media culture on a biplate of Middlebrook 7H10 agar with and without antibiotics, and a broth culture (BACTEC 960, Becton Dickinson and Company, Sparks, MD, VersaTrek, Thermofisher, formerly Trek Diagnostic Systems, Cleveland, Ohio) as previously described (18). *M. intracullulare* isolates were identified using AccuProbe (Hologic-GenProbe, San Diego, CA, as previously described (18). *In vitro* susceptibility testing of MAC isolates was performed as previously described (17). *M. intracellulare* growth on broth and solid media was assessed using semi-quantitative scoring: growth on broth medium only = “pos”, growth in broth medium plus 1-49 colonies on solid medium = countable colonies (cc), 50-99 colonies on solid medium = 1+, 100-199 colonies on solid medium = 2+, 200-299 colonies on solid medium = 3+, greater than 300 colonies on solid medium = 4+ (19).

### Cytokine Analysis

Batched BAL supernatant and plasma samples were stored at -80°C. They were then collectively thawed and analyzed in duplicate using the Invitrogen Cytokine Monkey Magnetic 28-plex Panel which includes monocyte chemo-attractant protein 1 (MCP-1), interleukin-12 (IL-12), granulocyte-monocyte colony stimulating factor (GM-CSF), macrophage inflammatory protein 1 beta (MIP 1-β), interferon-gamma (IFNγ), monokine induced by interferon-gamma (MIG), migration inhibition factor (MIF), IL-1 receptor antagonist (IL-1Ra), tumor necrosis factor (TNF), IL-2, IL-4, IL-8, intercellular adhesion molecule (ICAM), and RANTES (Regulated on Activation, normal T cell Expressed and Secreted). For animals sacrificed at day 30, plasma samples were obtained at baseline and on days 7, 14, 21, 30 with BAL samples were collected at day 30. For animals sacrificed at day 60, plasma samples were collected at baseline and on days 7, 14, 21, 28, and 60 along with BAL samples collected at day 60.

### Tissue Analysis

At the time of sacrifice, fresh tissues from the mediastinal lymph nodes, liver, spleen, and kidneys were prepared for culture. The lungs were resected *en-bloc* and inspected visually for areas of inflammation. One lobe that appeared abnormal was isolated, tied off, and resected for culture. The remaining segments of lung were suspended after cannulating the trachea and then inflated with formalin infused at a height of 30 centimeters. After inflation, the tracheal was tied off and the lungs; as well as sections from the liver, spleen, and kidneys were submerged in 100% formalin for a period of 10 days. All tissues were then sent to the Central Pathology Lab at the UTHSC-SA for processing and staining.

### Statistical analysis

While we reported for the variability of the cytokines, mean ± SD, statistical analyses were not performed because of the few number of animals used. We believe the trends of several of the cytokines between the *M. intracellulare*-inoculated marmosets that resulted in culturable *M. intracellulare* vs. non-culturable *M. intracellulare*, as well as the those with culturable *M. intracellulare* over the course of the infection, are compelling in this pilot, descriptive paper

## RESULTS

### Clinical and standard laboratory assessments of the *M. intracellulare*-infected marmosets

All animals had normal growth and behavior as well as normal blood chemistries and cellular counts prior to infection with *M. intracellulare*. Following instillation of *M. intracellulare*, the animals were regularly assessed for evidence of illness as per protocol for Texas Biomed. Throughout the study period, none of the animals displayed signs of respiratory disease such as cough or tachypnea, or weight loss (Table 1). All the marmosets exhibited normal behavior and activity until the time of sacrifice.

**TABLE 1:** Summary of Microbiologic, Radiographic and Pathologic Findings From Each Animal.

| Animal<br>(wt-pre/post, gms) | Day<br>sacrificed | *M. intracellulare* AFB culture<sup>1,2</sup> |  |  |  |  | CXR | Path |
| --- | --- | --- | --- | --- | --- | --- | --- | --- |
|  |  | Lung | Spleen | Kidney | Liver | BAL |  |  |
| 1<br>(474/473) | 30 | cc | cc | cc | — | — | + | + |
| 2<br>(390/388) | 30 | 3+ | cc | — | — | — | + | + |
| 3<br>(480/481) | 30 | —<br>- | —<br>- | —<br>- | —<br>- | — | — | — |
| 4<br>(500/497) | 60 | cc | — | — | — | — | + | + |
| 5<br>(401/408) | 60 | cc | cc | — | pos | — | + | + |
| 6<br>(486/472) | 60 | cc | — | —<br>- | —<br>- | — | + | + |
| 7<br>(475/473) | 60 | —<br>- | —<br>- | —<br>- | —<br>- | — | — | — |
| **Chest X-ray<sup>3</sup>**<br>1: RLL posterior consolidation<br>2: RLL posterior consolidation<br>3: Unremarkable<br>4: RLL posterior consolidation<br>5: RLL posterior consolidation<br>6: RLL posterior consolidation<br>7: Unremarkable |  | **Lung histopathology<sup>3</sup>**<br>1: RML, RLL, Left lung: diffuse involvement with lymphs & monocytes. Early granulomatous inflammation, few giant cells. Subpleural and mediastinal subscapular inflammation<br>2: Right lung 60% consolidated, mixed lymphs, monocytes, numerous granulomas and giant cells, significant edema around lymphatics. Left lung 10% consolidation, similar findings<br>3: RML, RLL, LLL: normal<br>4: RUL, RLL, LLL: Resolving inflammation with lymphs /monocytes - worse in subpleural region; no granulomas; RML airway damage extending to the pleural surface with bronchiectasis<br>5: RUL, RLL, LLL: Mild/moderate inflammation with lymphs/ monocytes, and some PMNs RLL<br>6: RUL, RLL, LLL: Mild/moderate inflammation with lymphs/ monocytes, RLL with early bronchiectasis<br>7: RML, RLL, LLL: Normal |  |  |  |  |  |  |

<sup>1</sup> *M. intracellulare* growth on broth and solid media: growth on broth medium only = “pos”, growth in broth medium plus 1-49 colonies on solid medium = countable colonies (cc), 50-99
colonies on solid medium = 1+, 100-199 colonies on solid medium = 2+, 200-299 colonies on solid medium = 3+, greater than 300 colonies on solid medium = 4+ (19).
<sup>2</sup>No specimens submitted for AFB stain (smear) and culture to the mycobacteriology laboratory had a positive AFB stain although some lung specimens submitted for histopathologic analysis were AFB stain (smear) positive. (Figure 7)
<sup>3</sup>RUL=right upper lobe, RML=right middle lobe, RLL=right lower lobe, LUL=left upper lobe, LLL=left lower lobe

### Mycobacterial burden

For the three marmosets that were sacrificed at day 30, two were culture positive for *M. intracellulare* in the lungs and spleens, and one was culture negative for the mycobacteria in all organs (Table 1). Of the two animals with lung culture positivity, one also had positive *M. intracellulare* cultures in the spleen and kidneys and the other also had positive culture in the spleen. Interestingly, all three animals had negative BAL cultures at day 30 just prior to euthanasia. No specimen submitted for AFB stain (smear) and culture was AFB stain positive.

For the four marmosets that were sacrificed at day 60, three had positive *M. intracellulare* cultures of the lungs and one was negative for the mycobacteria in all organs (Table 1). Of the three animals with positive *M. intracellulare* lung cultures, only one had a positive extra-pulmonary culture in the spleen and liver. Similar to the marmosets infected for 30 days, all had negative BAL cultures for mycobacteria at day 60. No specimen submitted for AFB culture was AFB stain positive, although some tissue specimens were AFB smear positive on histopathologic analysis.

On semi-quantitative analysis, all positive AFB cultures were scored “countable colonies”, 1-49 colonies on solid media) with two exceptions (Table 1). There was no apparent correlation between the degree of culture positivity from the lungs and the presence of positive extra-pulmonary cultures. All positive *M. intracellulare* cultures had the identical *in vitro* susceptibility pattern as the originally instilled *M. intracellulare* isolate with macrolide and amikacin resistance. Staining with fluorochrome confirmed the presence of AFB in the *M. intracellulare*-infected lung tissue samples (Figure 1).

**Figure 1:**
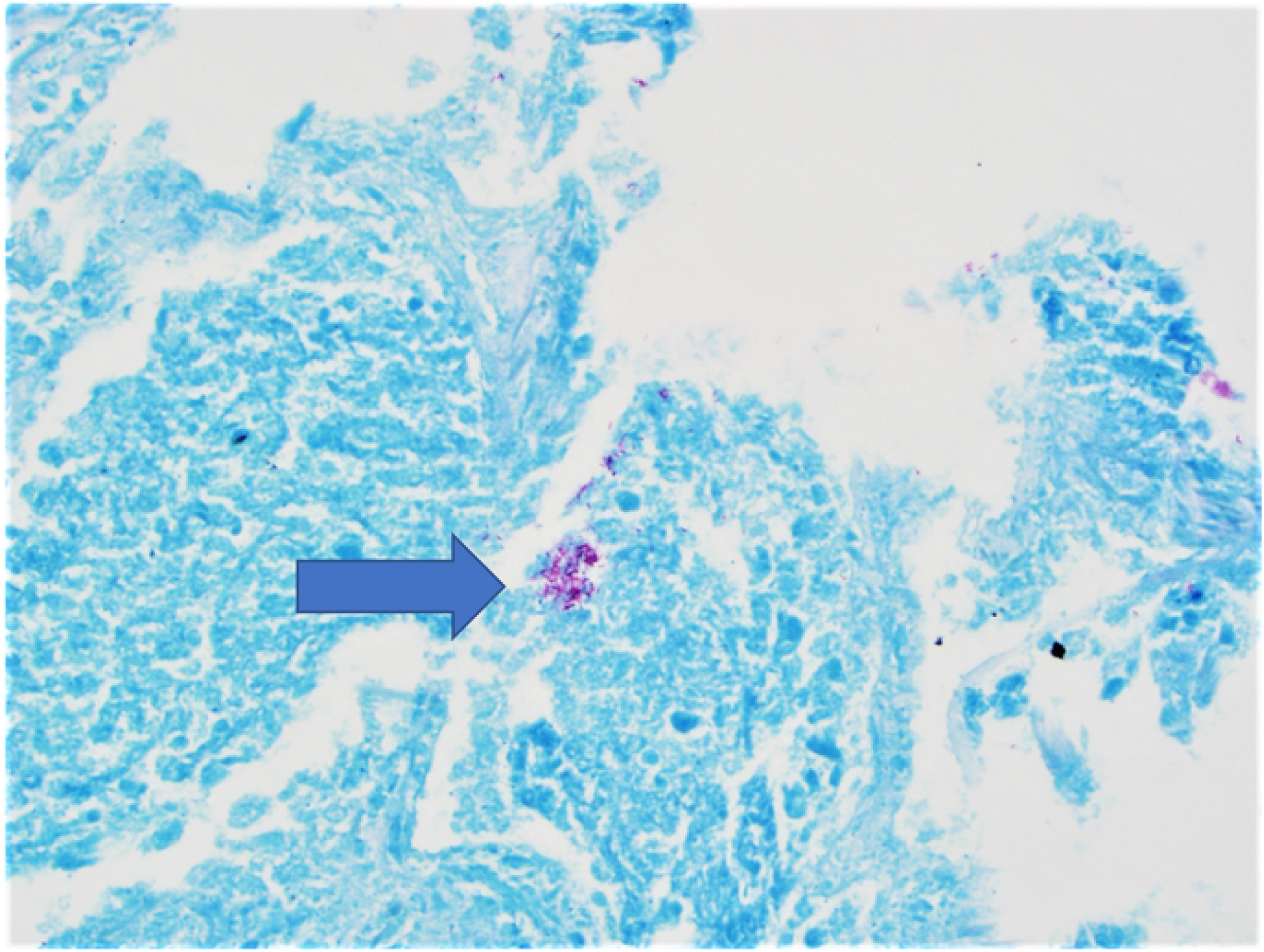
AFB detection. Positive AFB staining of MAC infected lung tissues on histopathologic examination from marmoset 2, sacrificed 30 days after inoculation with *M. intracellulare*.

### Chest radiographic features

The chest radiographs of all seven marmosets were within normal limits prior to infection with *M. intracellulare*. Following infection, the chest radiographs were abnormal with evidence of patchy consolidation in 5/5 animals with lung cultures positive for *M. intracellulare* (Figure 2). The animal with the culture score of “5” on semi-quantitative analysis also had the most extensive radiographic abnormalities (Table 1, Figure 1). The two animals with negative lung cultures for *M. intracelluare* had no end of study chest radiographic abnormalities.

**Figure 2:**
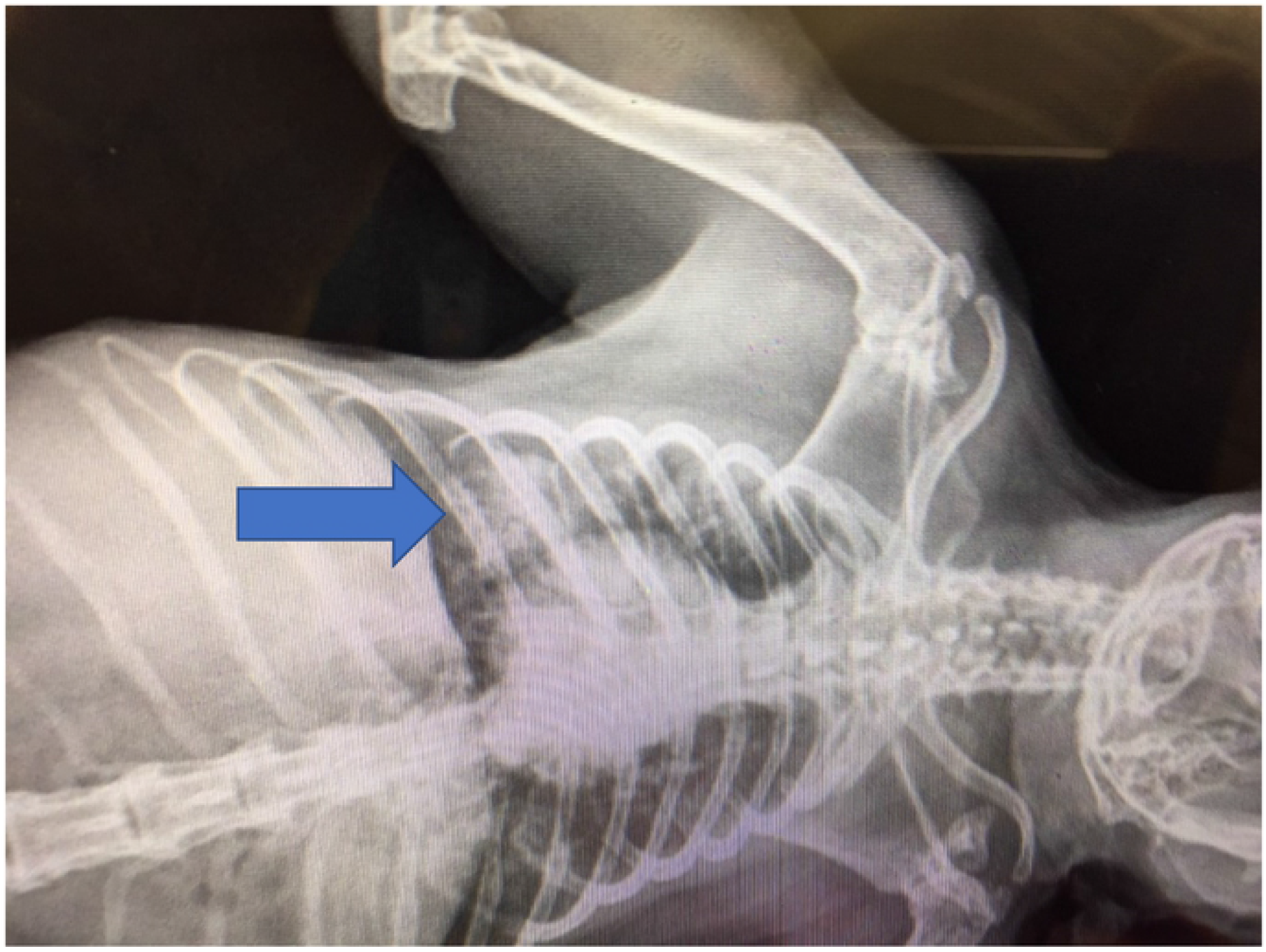
Chest radiograph prior to necropsy from Marmoset #2 showing right-sided density consistent with pneumonitis.

### Histopathologic findings

For the 5 infected animals, all five had visible hemorrhagic abnormalities in the lungs. Histopathologic features in the lungs of animals with culturable *M. intracelluare* at day 30 included mixed monocytic and lymphocytic infiltration around the bronchovascular bundle extending into the alveolar space. Numerous early granulomas with associated giant cells and occasional neutrophils and eosinophils were observed (Figure 3). Subpleural intra-alveolar nodules with severe pleuritis were seen in three animals. One animal had marked enlargement of the mediastinal lymph nodes with striking subcapsular mediastinal lymphadenitis. At day 60, animals were noted to have resolving inflammation in the lung parenchyma with residual bronchitis and bronchiolitis. Two animals had findings consistent with early bronchiectasis. Minimal or no granulomatous inflammation was noted by day 60. The spleen, kidney and liver histologic sections from the animals with positive cultures from those organs did not show abnormalities including evidence of inflammation typical of mycobacterial infection

**Figure 3:**
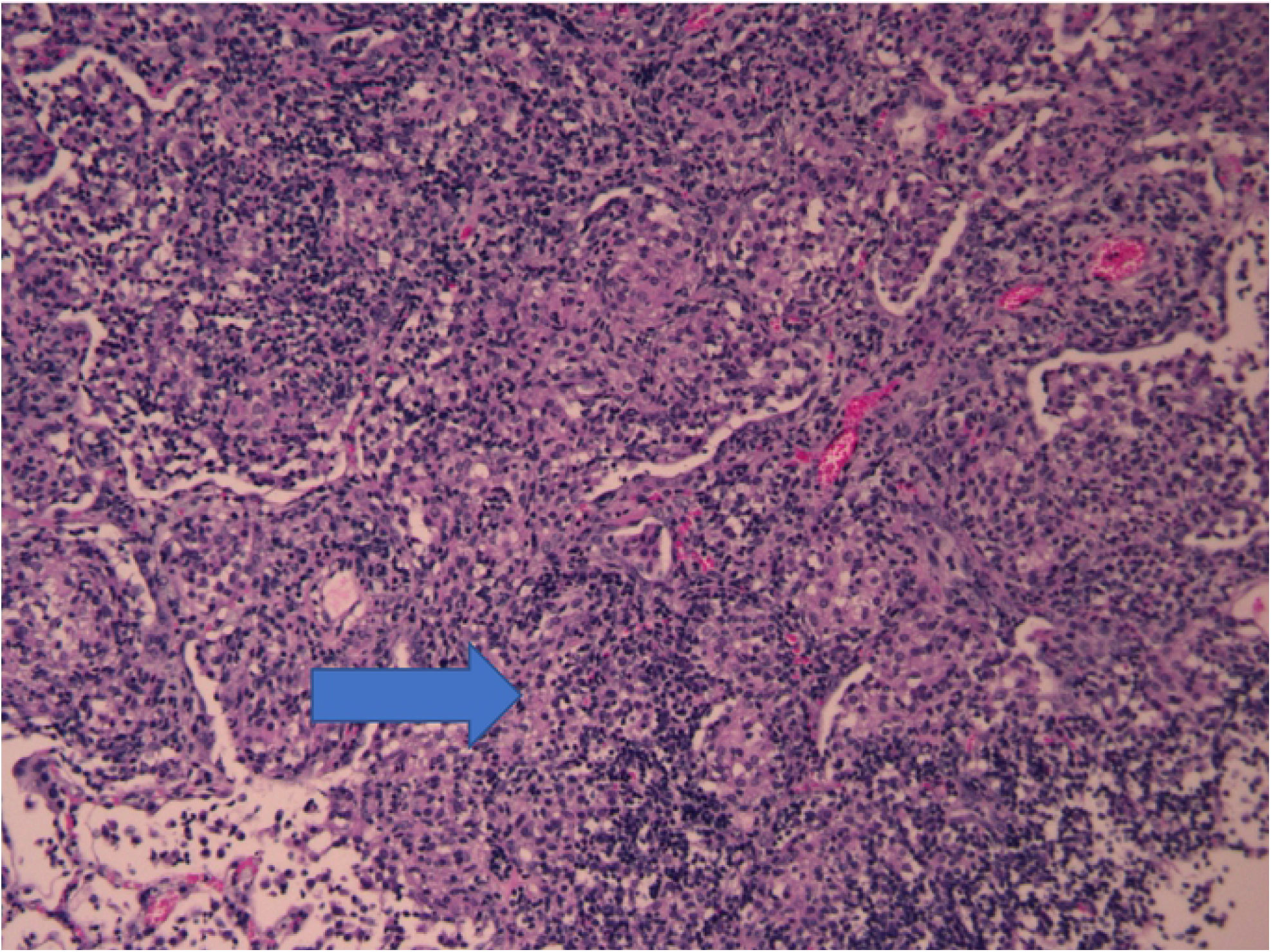
Animal #2, Day 30 RML showing early granuloma associated with giant cells and mixed lymphocytes/monocytes and occasional neutrophils and eosinophils.

### Serum cytokines and chemokines

Serum cytokine and chemokine levels were measured temporally from the time of infection to the time of sacrifice at day 30 or day 60. In the three animals sacrificed at day 30, the two with positive cultures for *M. intracellulare* had, in general, greater levels of MIF, MIP-1α, MIP-1β, IL-1Ra, IFNγ, and MIG than the lone animal with negative cultures (Figure 4A). Similarly, for the marmosets sacrificed at day 60, the same serum cytokine and chemokine levels for the three with a productive infection were generally greater when compared to the one animal with negative cultures (Figure 4B). For both the 30 and 60 days of infection, the cytokine and chemokine levels peaked approximately 14-30 days post-infection.

**Figure 4A:**
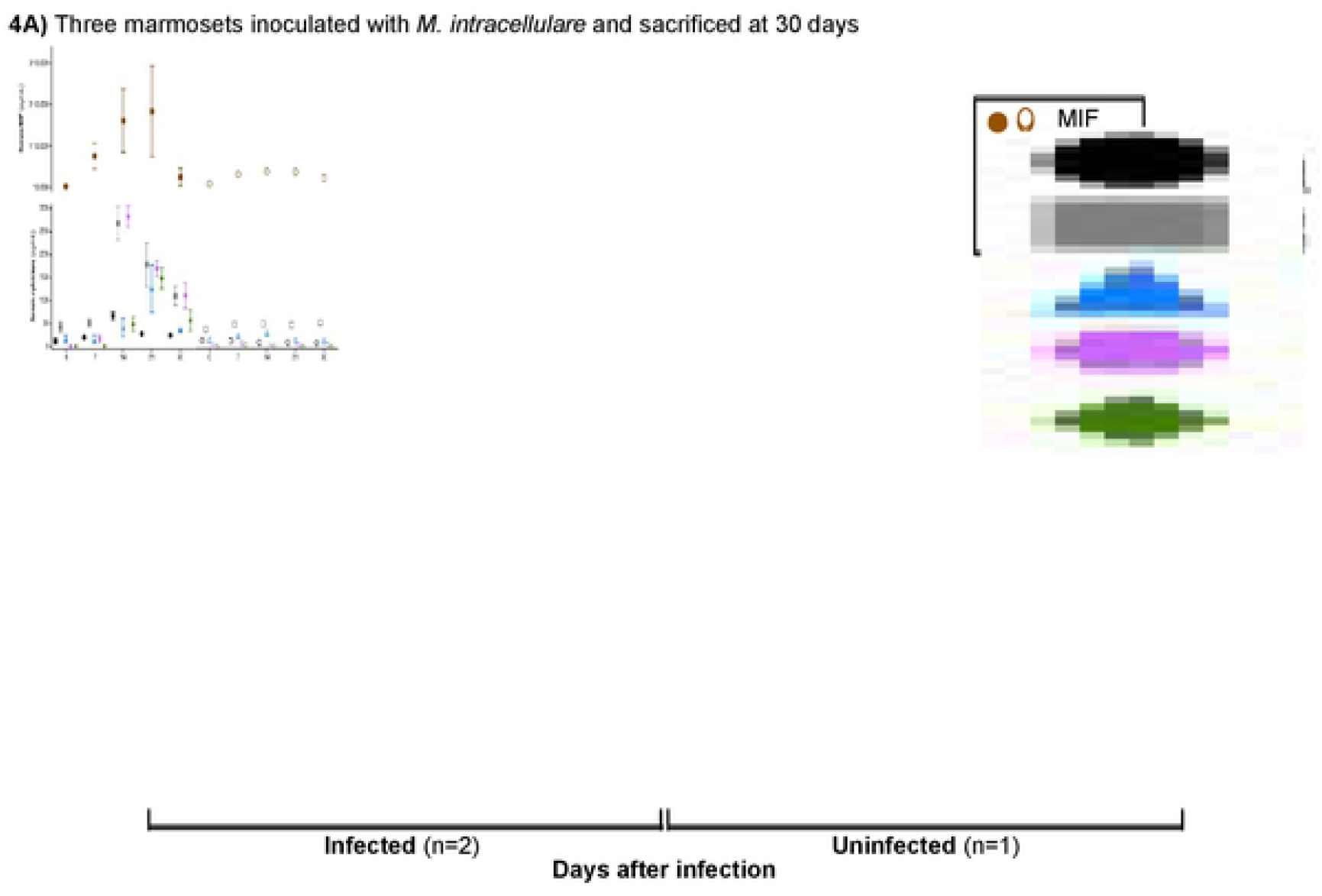
Serum cytokine/chemokine detection from day 0 to day 30: Sera from two animals with positive *M*. *intracellulare* cultures and one animal with negative *M. intracellulare* cultures were evaluated for various cytokine/chemokine expressions at indicated time points until day 30.

**Figure 4B:**
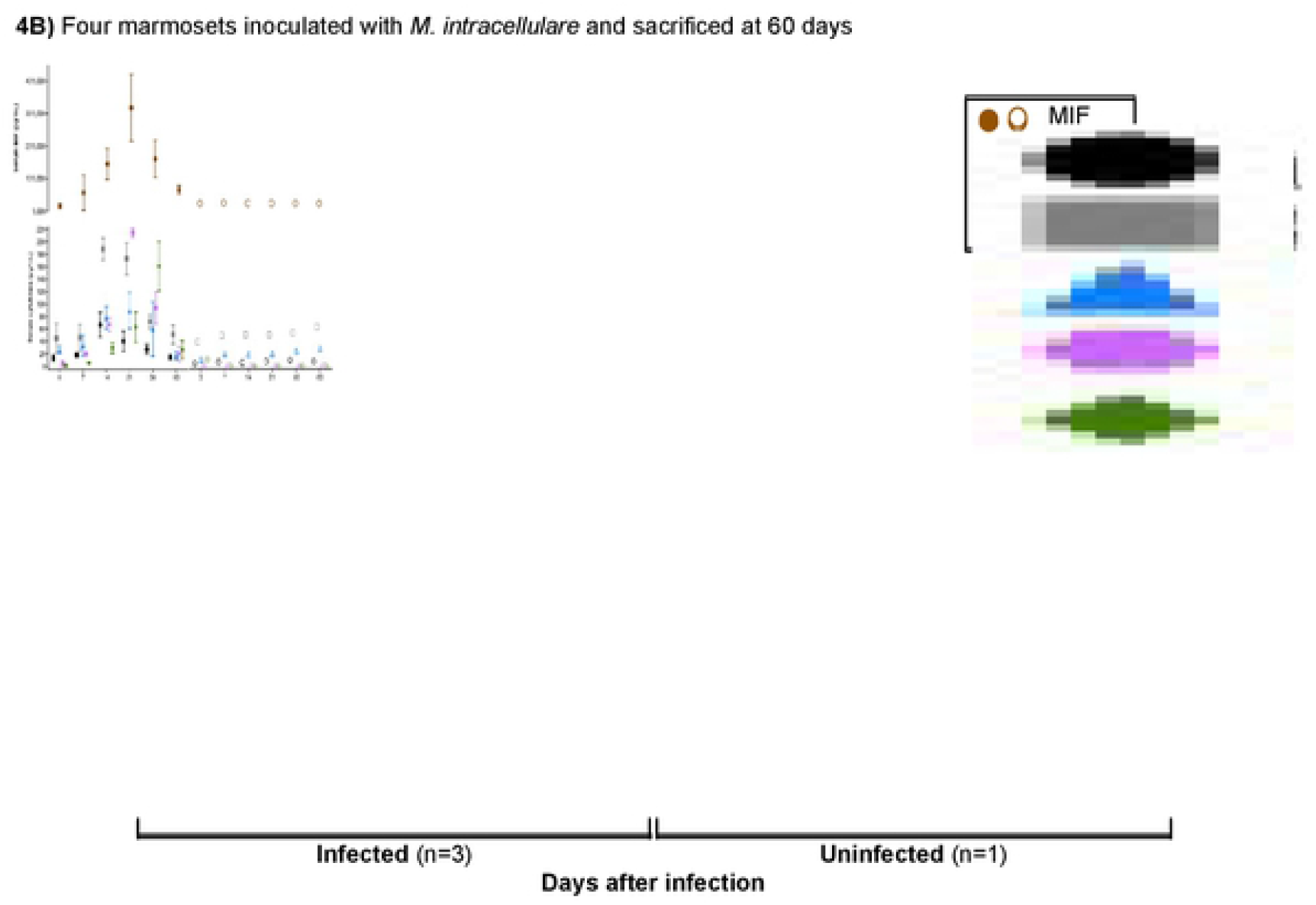
Serum cytokine/chemokine detection from day 0 to day 60: Sera from three animals with positive *M. intracellulare* lung cultures and one with negative lung culture were evaluated for various cytokine expressions at indicated time points untilday 60. These animals are different than the animals presented in Figure 4A.

### BAL cytokines

Cytokine and chemokine levels were also quantified in the BAL fluids of all animals prior to sacrifice at either day 30 or day 60. To varying degrees, BAL cytokine levels from the animals with positive lung cultures at either the day 30 or day 60 of infection (total n = 5) had increased levels of MIF, MIP-1α, MIP-1β, IL-1Ra, MIG, ICAM, IFNγ, RANTES, and TNF, compared to animals without a productive infection (total n = 2) (Figure 5A/5B). BAL levels for all the cytokines and chemokines were greater at day 30 except for MIF, which was greater at day 60.

**Figure 5A and 5B.**
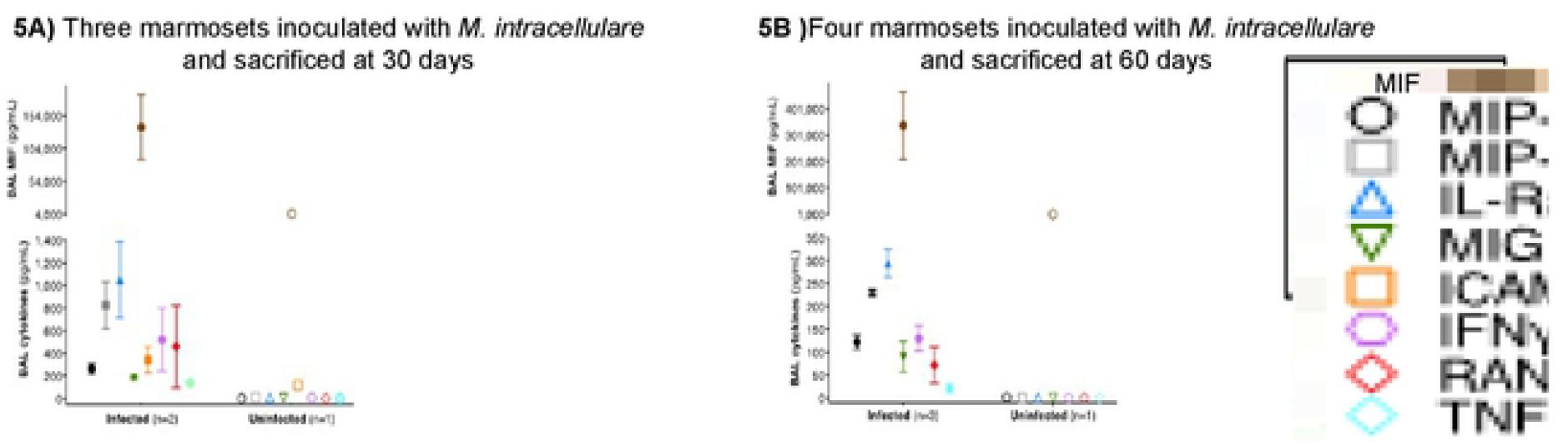
Cytokine detection in BAL from marmosets. A) At the time of sacrifice at day 30 after inoculation, the BAL fluids from two marmosets with *M. intracellulare-positive* lung cultures and one animal with negative cultures were analyzed for various cytokine/chemokine concentrations. B) At the time of sacrifice at day 60 after inoculation, the BAL fluids from three marmosets with *M. intrace/Julare-positive* lung cultures and one animal with negative cultures were analyzed for various cytokine and chemokine concentrations. These animals are different than the animals presented in Figure 5A.

Cytokines and chemokines in the 28-cytokine kit not mentioned were below the detection limit for both the serum and BAL assays.

## DISCUSSION

In this study, we successfully established pulmonary *M. intracellulare* infection in five of the seven healthy female marmosets. We chose female marmosets for this investigation due to the predominance of human *M. avium* and *M. intracellulare* lung disease in women (3-5). The animals showed no discernable clinical signs of lung infection, such as cough, decreased activity or weight loss, suggesting that the inoculation resulted in a “sub-clinical” chronic infection. While there was culture positivity in extra-pulmonary organs in a few animals, there were no accompanying histopathologic findings. This type of indolent infection is consistent with the nodular/bronchiectasis form of *M. intracellulare* lung disease in humans (3-5).

All *M. intracellulare* isolates cultured from the infected animals showed the same *in vitro* susceptibility pattern as the *M. intracellulare* isolate instilled in the animals, specifically, *in vitro* resistance to macrolide and amikacin, precluding the possibility of environmental *M. intracellulare* contamination of the tissue (and BAL) specimens. Two animals, one sacrificed at 30 days and one at 60 days, did not show evidence of infection. For both of these animals, there was *a priori* a question about the adequacy of the endotracheal *M. intracellulare* challenge.

The serum and BAL cytokine and chemokine levels for animals sacrificed at 30 and 60 days showed variability between animals but there were consistently higher levels for five animals with a productive *M. intracellulare* infection compared to the two animals without recoverable *M. intracelluare* from any of the organs.

Comparing serum and BAL cytokine and chemokine levels at both 30 days and 60 days for each cytokine, two general findings are that the serum levels peaked approximately day 14-30 that tapered off by day 60 whereas the BAL cytokine and chemokine levels remained elevated in the BAL even at day 60 in the *M. intracellulare* infected animals, suggesting a stronger local effect of the infection than a systemic one. Although there were consistent trends in cytokine/chemokine response to infection there was considerable variability between animals in the magnitude of response.

Our findings are consistent with those published for other mycobacterial diseases (20). MIP-1 α/β (now known as CCL3/CCL4) – members of the C-C superfamily associated with the early immune response – peaked at day 14 in the *M. intracellulare*-infected animals. This may account for the mixed monocytic/neutrophilic response seen in our model. IFNγ, considered to be essential for the induction of granulomatous inflammation, peaked between days 14-21. This was associated with a rise in MIF, known to be induced by IFNγ, which peaked between days 14-30. MIF is felt to contribute to detrimental inflammation but may be crucial in controlling infection caused by mycobacterial diseases (21,22). RANTES (CCL-5), which promotes macrophage chemotaxis and upregulation, peaked between days 30-60. High levels of TNF, the only serum cytokine not persistently higher at day 60, has been associated with necrosis in animal models of tuberculosis and was not observed in our model. As noted, although there were consistent trends in cytokine/chemokine response to infection in our study, there was considerable variability between animals in the magnitude of response.

Histopathological analysis of the *M. intracellulare* culture-positive lungs showed typical abnormalities consistent with granulomatous inflammation with giant cells but no evidence of necrosis, the latter consistent with the absence of TNF. Stains of lung tissue were inconsistently AFB smear positive (Figure 1) yet consistently culture positive for *M. intracellulare*, similar to findings in patients with MAC lung disease (3-5). Additionally, by day 60, the lungs showed resolving inflammation with persistent bronchitis and bronchiolitis and early evidence of bronchiectasis. These latter findings, in the context of persistently positive MAC cultures, would be consistent with chronic mycobacterial lung infections in humans.

Fibro-cavitary forms of MAC lung infection, including *M. intracellulare*, are clearly associated with bronchiectasis formation but it has never been established in the nodular/bronchiectasis form of MAC lung disease whether MAC can initiate bronchiectasis formation (23). The current consensus is that bronchiectasis likely precedes MAC infection for most patients and is, in fact, the critical predisposition for the establishment of MAC lung infection (3-5). Our results suggest that the MAC infection per se has the potential to cause bronchiectasis formation. Another possibility is that if some degree of pre-existing bronchiectasis is present prior to the establishment of MAC lung infection the bronchiectasis may contribute to or accelerate further bronchiectasis formation.

Winthrop et al. published results with three rhesus macaques infected with escalating doses (10^7^, 10^8^, 10^9^ CFU) of *M. avium* subsp. *hominissuis* strain 101 administered endobronchially (10). The animals that received 10^9^ CFU *M. avium* developed a right lung opacity radiographically on day 14 post-infection. The radiographic abnormality was associated with recovery of *M. avium* in BAL fluid culture. Similar to our findings, there were mid-infection increases in circulating cytokines, IL-6, IL-12 and IFNγ with peak levels at day 14 post-inoculation for IL-12, day 21 for IFNγ and day 28 for IL-6. BAL cytokines that were elevated peaked at mid-infection (42 days post inoculation) included, IL-6, IL-12, IFNγ, TNF, MIP-1β. Interestingly, only IL-6 did not return to baseline at the end of the infection period.

Our findings, from a pathophysiologic perspective, are generally consistent with the findings from Winthrop et al (10). It is possible that some dissimilarities could be due to differences in virulence and pathogenicity between *M. intracellulare* and *M. avium* (as has been noted in humans), differences in disease pathophysiology between the rhesus macaque and marmoset, or both (6,7).

A novel finding in this animal model was the positive cultures for *M. intracellulare* in extra-pulmonary organs including spleen, liver and kidney, without apparent clinically detectable infection in the extra-pulmonary organs as well as an absence of discernable inflammatory response in the involved organs. It is possible that the apparent dissemination of the organism is related to the relative susceptibility of marmosets to mycobacteria, although in contrast to TB disease in marmosets, the *M. intracellulare* infected animals did not die or even develop clinical signs of disease with the dissemination. Alternatively, *M. intracellulare* lung infection may be pathophysiologically similarly to a latent TB state with early dissemination followed by immune-mediated control of the infection in extra-pulmonary organs. The emergence of disseminated *M. avium* complex disease in individuals with advanced AIDS would lend credence to this notion of a latent NTM infection with subsequent reactivation with development of an immunocompromised state.

We cannot confidently assert that any current animal model faithfully replicates the pathophysiologic events associated with human MAC lung disease. With the current level of knowledge, there is no reliable way to correlate the animal findings of early and evolving MAC infection with those from humans, which are largely unknown. While this study does not present a comprehensive description of MAC lung disease pathophysiology, we believe that potential exists with the marmoset MAC lung disease model to accomplish this goal.

One important potential advantage of marmosets for studying *M. intracellulare* or *M. avium* lung disease is that they are relatively susceptible to NTM lung infection so that establishment of *M. intracellulare* lung infection does not require an immune compromised state or airway injury. Based on our success in establishing *M. intracellulare* infection in 5/7 challenged animals, marmosets appear to be a reproducible model for establishing *M. intracellulare* lung infection which could impact several areas of investigation. First, this model could more rigorously identify mechanisms of host susceptibility and disease progression pertinent to humans. For example, marmosets could help identify virulence differences between *M. intracellulare* and *M. avium* and might also serve as a model for less common NTM pathogens such as *M. abscessus* or *M. xenopi*. Second, a reliable non-human primate model of MAC lung infection could also prove more informative and predictive of drug responses than current non-primate models. Third, the development of bronchiectasis in the marmosets, even after a short follow-up time, could provide insights into the pathophysiology of this complex process. Fourth, translational studies comparing the immune response in the lung tissues of MAC-infected marmosets with that of surgically removed lung tissues of MAC lung disease patients may provide insights into the pathogenesis of progressive NTM lung disease in humans.

There are several limitations to this study. We did not assess cellular activity prior to and after challenge with *M. intracellulare*. Chest CT and PET-CT were not obtained and would have been more sensitive for exposing areas of mycobacterial infection.

In conclusion, we demonstrated that endobronchial instillation of *M. intracelluare* reliably results in pulmonary mycobacterial infection in marmosets. The infection is associated with a reproducible immune reaction, radiographic abnormalities and histologic changes consistent with mycobacterial lung infection in humans. Additionally, the infection is indolent without visible acute harm or impact to the animal, similar to the *M. intracellulare*-associated lung disease in humans. We believe this model has the potential to answer questions about *M. intracellulare* disease pathophysiology as well as the natural history of *M. intracellulare* lung disease. Future investigations will include longer post-exposure observation of the animals and more extensive pathophysiologic investigation.

## Acknowledgments

We are grateful to Lore Fornis for expert help with the graphs and to Dr. Matthew Strand in the Biostatistics Department at National Jewish Health for intellectual input.

## Notes

Financial Support: Harry L. Willett Foundation, Denver, CO.

### Competing Interest Statement

The authors have declared no competing interest.

## References

1: Daniel-Wayman S, Adjemian J, Prevots DR. Epidemiology of Nontuberculous Mycobacterial Pulmonary Disease (NTM PD) in the USA. In: Griffith DE (Ed), Nontuberculous Mycobacterial Disease: a comprehensive approach to diagnosis and management, 1st Edition, Springer Nature Switzerland AG, Cham, Switzerland; 2019. Pg 145–162.

2: Wagner D, Lipman M, Cooray S, Ringshausen FC, Morimoto K, Koh W-J, Thomson R. Global Epidemiology of NTM Disease (Except North America). In: Griffith DE (Ed), Nontuberculous Mycobacterial Disease: a comprehensive approach to diagnosis and management, 1st Edition, Springer Nature Switzerland AG, Cham, Switzerland; 2019. Pg 163–260.

3: Holt MR, Daley CL. Mycobacterium avium complex Disease. In: Griffith DE (Ed), Nontuberculous Mycobacterial Disease: a comprehensive approach to diagnosis and management, 1st Edition, Springer Nature Switzerland AG, Cham, Switzerland; 2019. Pg 310–324.

4: Cowman S, van Ingen J, Griffith DE, Loebinger MR. Non-tuberculous mycobacterial pulmonary disease. Eur Respir J. 2019 Jul 11;54(1).

5: Griffith DE. Treatment of Mycobacterium avium Complex (MAC). Semin Respir Crit Care Med. 2018 Jun;39(3):351–361. doi: 10.1055/s-0038-1660472. Epub 2018 Aug 2.

6: Koh WJ, Jeong BH, Jeon K, Lee NY, Lee KS, Woo SY, Shin SJ, Kwon OJ. Clinical significance of the differentiation between Mycobacterium avium and Mycobacterium intracellulare in M. avium complex lung disease. Chest. 2012 Dec;142(6):1482–1488.

7: Kim SY, Shin SH, Moon SM, Yang B, Kim H, Kwon OJ, Huh HJ, Ki CS, Lee NY, Shin SJ, Koh WJ. Distribution and clinical significance of Mycobacterium avium complex species isolated from respiratory specimens. Diagn Microbiol Infect Dis. 2017 Jun;88(2):125–137.

8: Chan E. Vulnerability to Nontuberculous Mycobacterial Lung Disease or Systemic Infection Due to Genetic/Heritable Disorders. In: Griffith DE (Ed), Nontuberculous Mycobacterial Disease: a comprehensive approach to diagnosis and management, 1st Edition, Springer Nature Switzerland AG, Cham, Switzerland; 2019. Pg 89–110.

9: Blanchard JD, Elias V, Cipolla D, Gonda I, Bermudez LE. Effective Treatment of Mycobacterium avium subsp. hominissuis and Mycobacterium abscessus Species Infections in Macrophages, Biofilm, and Mice by Using Liposomal Ciprofloxacin. Antimicrob Agents Chemother. 2018 Sep 24;62(10).

10: Winthrop K, Rivera A, Engelmann F, Rose S, Lewis A, Ku J, Bermudez L, Messaoudi I. A Rhesus Macaque Model of Pulmonary Nontuberculous Mycobacterial Disease. Am J Respir Cell Mol Biol. 2016 Feb;54(2):170–6.

11: Via LE, Weiner DM, Schimel D, Lin PL, Dayao E, Tankersley SL, Cai Y, Coleman MT, Tomko J, Paripati P, Orandle M, Kastenmayer RJ, Tartakovsky M, Rosenthal A, Portevin D, Eum SY, Lahouar S, Gagneux S, Young DB, Flynn JL, Barry CE 3rd. Differential virulence and disease progression following Mycobacterium tuberculosis complex infection of the common marmoset (Callithrix jacchus). Infect Immun. 2013 Aug;81(8):2909–19.

12: Via LE, England K, Weiner DM, Schimel D, Zimmerman MD, Dayao E, Chen RY, Dodd LE, Richardson M, Robbins KK, Cai Y, Hammoud D, Herscovitch P, Dartois V, Flynn JL, Barry CE 3rd. A sterilizing tuberculosis treatment regimen is associated with faster clearance of bacteria in cavitary lesions in marmosets. Antimicrob Agents Chemother. 2015 Jul;59(7):4181–9.

13: Scanga CA, Flynn JL. Modeling tuberculosis in nonhuman primates. Cold Spring Harb Perspect Med. 2014 Sep 11;4(12): a018564.

14: Cadena AM, Klein EC, White AG, Tomko JA, Chedrick CL, Reed DS, Via LE, Lin PL, Flynn JL. Very Low Doses of Mycobacterium tuberculosis Yield Diverse Host Outcomes in Common Marmosets (Callithrix jacchus). Comp Med. 2016;66(5):412–419.

15: Weatherall D. The use of non-human primates in research. 2006. Available from, http://www.royalsoc.ac.uk/weatherall.

16: Guide for the Care and Use of Laboratory Animals of the National Institutes of Health (NIH), the Office of Animal Welfare, 8th ED. National Academies Press, Washington DC; 2011.

17: CLSI M24, Susceptibility Testing of Mycobacteria, Nocardia spp, and Other Aerobic Actinomycetes. 3rd Ed. Clinical Laboratory Standards Institute, Wayne, PA: 2018)

18: CLSI M48, Laboratory Detection and Identification of Mycobacteria. 2nd Ed. Clinical Laboratory Standards Institute, Wayne, PA: 2018)

19: Griffith DE, Adjemian J, Brown-Elliott BA, Philley JV, Prevots DR, Gaston C, Olivier KN, Wallace RJ Jr. Semiquantitative Culture Analysis during Therapy for Mycobacterium avium Complex Lung Disease. Am J Respir Crit Care Med. 2015 Sep 15;192(6):754–60.

20: Barnes PF, Lu S, Abrams JS, Wang E, Yamamura M, Modlin RL. Cytokine production at the site of disease in human tuberculosis. Infect Immun. 1993, Aug;61(8):3482–9.

21: Das R, Koo MS, Kim BH, Jacob ST, Subbian S, Yao J, Leng L, Levy R, Murchison C, Burman WJ, Moore CC, Scheld WM, David JR, Kaplan G, MacMicking JD, Bucala R. Macrophage migration inhibitory factor (MIF) is a critical mediator of the innate immune response to Mycobacterium tuberculosis. Proc Natl Acad Sci U S A. 2013 Aug 6;110(32): E2997–3006.

22: Gerrit Grieb, Melanie Merk, Jürgen Bernhagen, Richard Bucala. Macrophage migration inhibitory factor (MIF): a promising biomarker. Drug News Perspect. 2010 May; 23(4): 257–264.

23: Griffith DE, Aksamit TR. Bronchiectasis and nontuberculous mycobacterial disease. Clin Chest Med. 2012 Jun;33(2):283–95.

